# Ezetimibe inhibits Dengue virus infection in Huh-7 cells by blocking the cholesterol transporter Niemann–Pick C1-like 1 receptor

**DOI:** 10.1101/382747

**Authors:** Juan Fidel Osuna-Ramos, José Manuel Reyes-Ruiz, Patricia Bautista-Carbajal, Noe Farfán-Morales, Margot Cervantes-Salazar, Rosa María del Ángel

## Abstract

Despite the importance of Dengue virus (DENV) infection in human health, there is not a fully effective vaccine or antiviral treatment against the infection. Since lipids such as cholesterol are required during DENV infection, its uptake and synthesis are increased in infected cells. Ezetimibe is an FDA-approved drug that reduces cholesterol uptake in humans by inhibiting the endocytosis through Niemman-Pick C1-Like 1 (NPC1L1) receptor, expressed on the membrane of enterocytes and hepatocytes. Our results indicate that an increase in the amount of NPC1L1 occurs on the surface of Huh-7 cells during DENV infection, which correlates with an increase in cholesterol levels. Blockage of NPC1L1 with ezetimibe in concentrations up to 50 μM does not reduce cell viability but diminished total cellular cholesterol, the percentage of infected cells, viral yield, viral RNA and protein synthesis without affecting DENV binding and/or entry to Huh-7 cells. Moreover, ezetimibe inhibited DENV replicative complex formation and lipid droplets accumulation. All these results indicate that ezetimibe is an excellent drug to inhibit DENV infection and confirm that cholesterol is a key target to inhibit viral infection.

## Introduction

Dengue is a systemic infection caused by dengue virus (DENV), a member of the *Flaviviridae* family and *Flavivirus* genus, that includes four serotypes (DENV 1 to DENV 4) (Guzman and Harris, 2015). Dengue is the most important mosquito-borne viral diseases causing more than 96 million infections per year with approximately 3.6 billion people at risk, affecting more than 100 countries around the world (Hasan et al., 2016; Simmons et al., 2012). DENV infection can be an asymptomatic illness or can produce a febrile illness (dengue fever, DF) which can lead to the severe disease complications, dengue hemorrhagic fever (DHF) or the dengue shock syndrome (DSS) (Hadinegoro, 2012). Currently, there are no specific drugs to treat the infection with DENV or a fully effective vaccine. Indeed, viral factors such as the high genetic variability among the four serotypes of DENV (Lim et al., 2013), antibody-dependent enhancement (ADE), and cross-reactivity with other *Flavivirus* such as Zika virus, make difficult to obtain a fully effective vaccine (Halstead and Russell, 2016; Villar et al., 2015). For this reason, development of therapeutic strategies with a pan-serotype effect is a priority. One possibility for the development of therapeutic strategies is to identify host factors that are usurped by DENV for replication (Acosta and Bartenschlager, 2016). A common feature among the members of the *Flaviviridae* family is the induction of cell membranes formation in the endoplasmic reticulum (ER) that forms part of structures, rich in lipids, known as replication complexes (RC) (Welsch et al., 2009). On the other hand, different lipids, such as cholesterol, are essential components of the virion (Carro and Damonte, 2013) and cholesterol is also required for RC formation (Rothwell et al., 2009a; Soto-Acosta et al., 2013). Thus, cholesterol could represent a key element at multiple stages of DENV replicative cycle. The cellular cholesterol content depends on the balance between uptake and *de novo* synthesis (Simons and Ikonen, 2000). Previous studies conducted by our group, reported an increase in the low-density lipoprotein receptor (LDLr) on the cell surface at early times after DENV infection (Soto-Acosta et al., 2013), suggesting that extracellular cholesterol is an essential source of this compound during DENV infection. Given the fact that several molecules are cholesterol receptors in the cell, we decided to study the participation of the Niemman-Pick C1-Like 1 (NPC1L1) receptor, which is a crucial molecule for the absorption and homeostasis of cholesterol in the human body (Betters and Yu, 2010a). This receptor is found on the cellular surface of enterocytes and hepatocytes and is responsible for the cellular absorption of cholesterol in hepatocytes (Jia et al., 2011). Some evidence suggests that the molecular mechanism for cholesterol absorption by the NPC1L1 receptor is dependent on clathrin-mediated endocytosis (Betters and Yu, 2010b). Interestingly, the internalization of the receptor can be blocked by ezetimibe, a novel FDA-approved drug used to lower serum cholesterol in patients with dyslipidemia, when other lipid-lowering drugs have failed (Betters and Yu, 2010b; Chang and Chang, 2008; Weinglass et al., 2008). Our results indicate that NPC1L1 receptor participates actively in the cholesterol uptake during DENV infection and that the drug ezetimibe inhibits DENV infection in the human hepatoma cell line (Huh-7) by blocking the NPC1L1 receptor.

## Materials and Methods

### Cell culture, virus, and reagents

The human hepatoma-derived cell line Huh-7 (Kindly donated by Dr. Ana Maria Rivas, Autonomous University of Nuevo León) and the Vero cells (Green monkey kidney cells, ATCC CCL-81) were grown in advanced DMEM (complete medium) supplemented with 2 mM glutamine, penicillin (5 x 10^4^) streptomycin (50 μg/mL), 8% fetal calf serum (FCS) and 1 mL/L of amphotericin B (Fungizone) at 37° C and a 5% CO_2_ atmosphere.

The propagation of DENV serotype 2 (New guinea strain) and DENV serotype 4 (H241 strain) was carried out using CD-1 suckling mice brains (provided by the Unidad de Producción y Experimentación de Animales de Laboratorio (UPEAL)). The Brain extracts from MOCK-infected CD-1 suckling mice were used as a control. Viral titers were determined by foci forming units assay in Huh-7 cells.

As reagents, anti-prM-E monoclonal 2H2 (ATCC^®^ HB-114) antibody, anti-E (Genetex), anti-NS3 (Genetex) and anti-C (Genetex) antibodies were used in flow cytometry and confocal microscopy to stain DENV viral proteins. The mouse anti-NPC1L1 (Santa Cruz Biotechnologies) polyclonal antibody was used to stain Niemann-Pick C1-Like 1 receptor on the cellular surface. The drug Ezetimibe was used for the inhibitions assays and was obtained from Cayman Chemical (Ann Arbor, MI).

### DENV infection and treatment

The Huh-7 cells seeded in 12-wells plate format at 70-80% of confluence were infected with DENV (serotype 2 or 4) at a multiplicity of infection (MOI) of 3 in medium supplemented with 1% of FCS for 2 hours (h) at 37°C. Later, cells were washed three times with HANKS’ solution and later the inhibition assay was performed using vehicle-treatment (Ethanol) or treatment with ezetimibe in complete medium for 48 h at 37°C.

### Cell viability assay and IC50 determination

The cell viability in cells treated with vehicle or with increasing concentrations of ezetimibe (0 μM (vehicle), 25 μM, and 50 μM ezetimibe) for 48 h at 37°C was assayed using propidium iodide (PI) and MTT methods. The cell viability was interpreted as the percentage of live cells by flow cytometry (BD LSR Fortessa™) by the PI method, while the MTT method was evaluated by spectrophotometry (BioTek ELx800) measuring the absorbance at 540 nm. The IC50 was calculated over a wider range of concentrations (5, 10, 15, 25, 45, 50, 60 μM ezetimibe) and the data was analyzed in the Graph Pad Prism software.

### Viral yield

Supernatants from infected untreated and treated cells were used to determine the viral yield using foci forming units (FFU) assay. Briefly, confluent monolayers of Huh-7 cells grown in 96-well plates were incubated with serial dilutions of supernatants from DENV-infected untreated or treated cells (final volume 50 μL) and incubated for 2 h at 37 °C to allow viral absorption. Then, the inoculum was removed, the cells were washed three times with HANKS’ solution, and 200 μL of the complete medium was added. The medium was removed 48 hours post-infection (hpi) and the cells were fixed with 1% formaldehyde, permeabilized for 20 min (0.1% saponin and 1% FBS, in 1X PBS), incubated with the 2H2 anti-prM-E monoclonal antibody for 2 h at room temperature (RT), and detected with the anti-mouse Alexa 488 (Life technologies) secondary antibody. Foci were quantified under the fluorescence microscopy and expressed as Foci Forming Units per milliliter (FFU)/mL)

### Quantification of the viral genome by qRT-PCR

The qRT-PCR was performed as previously described (Bautista-Carbajal et al., 2017). Briefly, the cDNA was obtained by RT-PCR using 1 μg of total RNA from each experimental condition, the enzyme ImpromII (Promega), and random primers (0.025 μg/μL) (Promega) with the following conditions: 25°C for 5 min, 42°C for 60 min, and 70°C for 15 min (Veriti Thermal Cycler, Applied Biosystems). The qPCR was performed with the SYBR Fast universal kit (Chukkapalli et al.), 5 μL of 2X Master Mix, 1 μL of cDNA, and the primers previously described in an Eco Illumina System with the following conditions: 2 min at 50°C, 2 min at 95°C, 40 cycles of 5 s at 95°C, and 30 s at 55°C.

### Western blot assay

Huh-7 cells were infected with DENV 2 or 4 at a MOI of 3 and treated with ezetimibe (0, 25 and 50 μM). Cell lysates were obtained and analyzed at 48 hpi to determine the presence of the NS3 protein as infection control. A total of 35 μg of protein extract, preheated at 95 °C for 10 min in the presence of 3% β-mercaptoethanol, were separated by electrophoresis in 10% SDS-PAGE and transferred to nitrocellulose membrane (Bio-Rad) and immunoblotting using rabbit polyclonal antibody directed against the NS3 protein (GENETEX). The proteins were detected by the use of Super Signal West Femto Chemiluminescent Substrate (Thermo Scientific), and densitometric analysis was performed using the ImageJ software (NIH) adjusting with the loading control α-GAPDH.

### Flow cytometry and confocal microscopy

The Huh-7 infected untreated and treated cells were analyzed by flow cytometry and confocal microscopy to determine the percentage of infected cells and viral or cellular proteins localization. The cells were grown in 24-wells plates for confocal microscopy and in 12-wells plates for flow cytometry. The harvested cells or slides were fixed with 1% formaldehyde, and were permeabilized for 20 min with permeabilizing solution and were incubated with anti-prM-E (2H2), anti-NS3, anti-C, anti-E or anti-NS4A antibodies for 2 h at RT; only the anti-NPC1L1 antibody was incubated overnight at 4°C, with 10% FCS, 3% of Bovine serum albumin (BSA) and 10 mM of glycine, without permeabilizing solution. As secondary antibodies, a goat anti-mouse Alexa Flour 488, a goat anti-mouse Alexa Flour 405 and a goat anti-rabbit Alexa Flour 555 were used (Life Technologies). To stain intracellular cholesterol dye filipin III complex (Sigma) was used, and the nuclei were counterstained with DAPI or propidium iodide (Life Technologies). The samples were observed in a Zeiss LSM700 laser confocal microscopy and images were analyzed with ZEN software v. 2010. Flow cytometry was performed in a BD LSR Fortessa^™,^ and the data were analyzed using the FlowJo v. 10 software

### Transmission electron microscopy

Mock-infected, infected untreated and ezetimibe treated Huh-7 cells were washed three times with PBS, fixed with 2.5% glutaraldehyde in 0.1 M sodium cacodylate buffer (pH 7.2) for 1 h at RT. Then, the cells were treated with 1% osmium tetroxide in 0.1 M sodium cacodylate buffer for 1 h at RT and dehydrated gradually by different concentrations of ethanol and propylene oxide and included in Poly/bed epoxy resin at 60 °C during 24 h. Thin sections (70 nm) were stained with uranyl acetate and lead citrate and visualized in a Jeol JEM-1011 transmission electron microscope.

### Cellular cholesterol quantification

The cholesterol content was quantified by fluorometric enzymatic assay using a kit (Cayman Chemical Company, Ann Arbor, MI, USA), following the manufacturer’s instructions. MOCK-infected or DENV-infected cells untreated or ezetimibe treated were lysed with lysis buffer (Chloroform, isopropanol, NP-40, 7:11:0.1). Fluorescence was detected using emission wavelengths of 585 nm in a microplate reader (Infinite^®^ 200 PRO).

### Virus binding assay

The Huh-7 cells in suspension were pre-treated with 50 μM of ezetimibe or anti-NPC1L1 antibody (diluted 1:100) during 2 h at 37 °C. Later, the cells were washed three times with PBS and were incubated with DENV 2 or DENV 4 at 4 °C for 30 min. Then, cells were fixed at RT for 20 min with 1% formaldehyde, blocked with 2% FBS, incubated with anti-E MAb antibody (Genetex) diluted 1:100, and detected with goat anti mouse-Alexa 405 antibody (Life Technologies). Finally, 10,000 events were quantified by flow cytometry (BD LSR Fortessa™) and the data were analyzed with the FlowJo v. 10 software.

### Statistics

For the statistical analysis, the Graph Pad Prism software version 6.0 was used. Numerical data were expressed with the value of the means and the standard deviation (SD). For inhibition assays, the foci tests and the qRT-PCR the Kruskal-Wallis nonparametric test was used for multiple comparisons with the Dunn post-Hoc test. For the percentage of infected cells in the flow cytometry, the Student’s t-test with the Welch correction was used. Statistical significance was established at the 95% level (p = <0.05).

## Ethics statement

This study was conducted in accordance with the Official Mexican Standard Guidelines for Production, Care and Use of Laboratory Animals (NOM-062-ZOO-1999) and the protocol, number 048–02, was approved by the Animal Care and Use Committee (CICUAL) at CINVESTAV-IPN, Mexico.

## Results

### The amount of NPC1L1 receptor on cell surface increases during DENV infection

Since the main role of the Niemann-Pick C1-Like 1 (NPC1L1) receptor is the cholesterol uptake, and cholesterol is essential for DENV infection, the first step was to determine if this receptor was present on the cell surface during DENV infection. Thus, non-permeabilizing uninfected and infected Huh-7 cells were incubated with the anti-NPC1L1 receptor antibody and analyzed by confocal microscopy and flow cytometry. As it can be observed, during the first hour postinfection (hpi) an increase in the amount of NPC1L1 receptor on the cell surface compared to the uninfected cells (MOCK-infected) was observed (Fig. 1 A and B). However, at 3 and 6 hpi an abrupt reduction in the fluorescence detected on the surface of DENV infected cells was detected, suggesting that the receptor was internalized (Fig. 1 A and B). However, at 12 hpi the receptor was located again on the surface of the infected cells (Fig. 1 A and B). These results suggest that DENV infection induces an increase of the NPC1L1 receptor on the cell surface during viral entry (at 1hpi) and during viral replication (at 12 hpi). The decrease in the amount of the receptor at 3 and 6 hpi could be related to the cholesterol uptake by internalization and recycling of the receptor (Ge et al., 2008).

**Figure 1.**
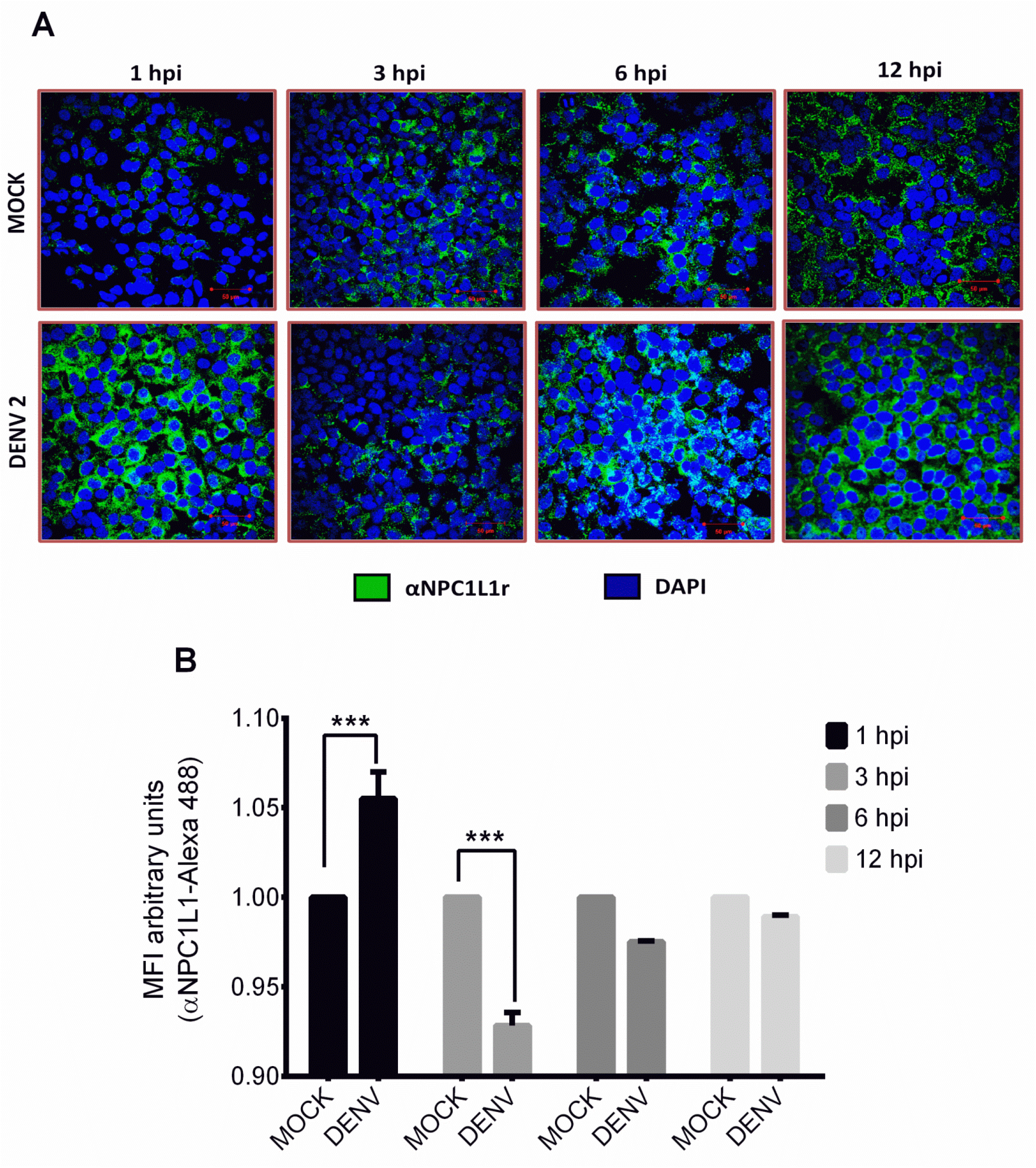
Localization of NPC1L1 receptor during DENV infection. Huh 7 cells were infected or not (MOCK) with DENV 2 at a MOI of 3, fixed at 1, 3, 6 and 12 hours post-infection (hpi) and the presence of NPC1L1 receptor (green) on the nonpermeabilized cells was analyzed by confocal microscopy (A) and flow cytometry (B). The nuclei of the cells were stained with DAPI (blue), The graphs represent the results expressed as MOCK-normalized mean and SEM of three independent experiments. \**p*= ≥ 0.0001.

### Ezetimibe reduces DENV infection

Once we determined that the amount of NPC1L1 receptor on the surface of Huh-7 cells increase early after infection, we decided to analyze its importance during DENV infection. To investigate this fact, cells were treated with different concentrations of ezetimibe or vehicle (0 μM). Although ezetimibe has not a negative effect on cell viability evaluated by IP and MTT methods (Fig. 2 A and B), it causes an arrest of the receptor on the cell membrane of infected cells at 1, 3, 6 and 12 hours post-infection, as could be observed by confocal microscopy from non-permeabilized cells (Fig. 2 C). Interestingly, ezetimibe was able to inhibit the amount of DENV 2 infected cells, evaluated by flow cytometry at 48 h postinfection with an IC50 value of 13.07 μM (95% Confidence Interval = 9.60 - 16.54 μM) (Fig. 3 A).

**Figure 2.**
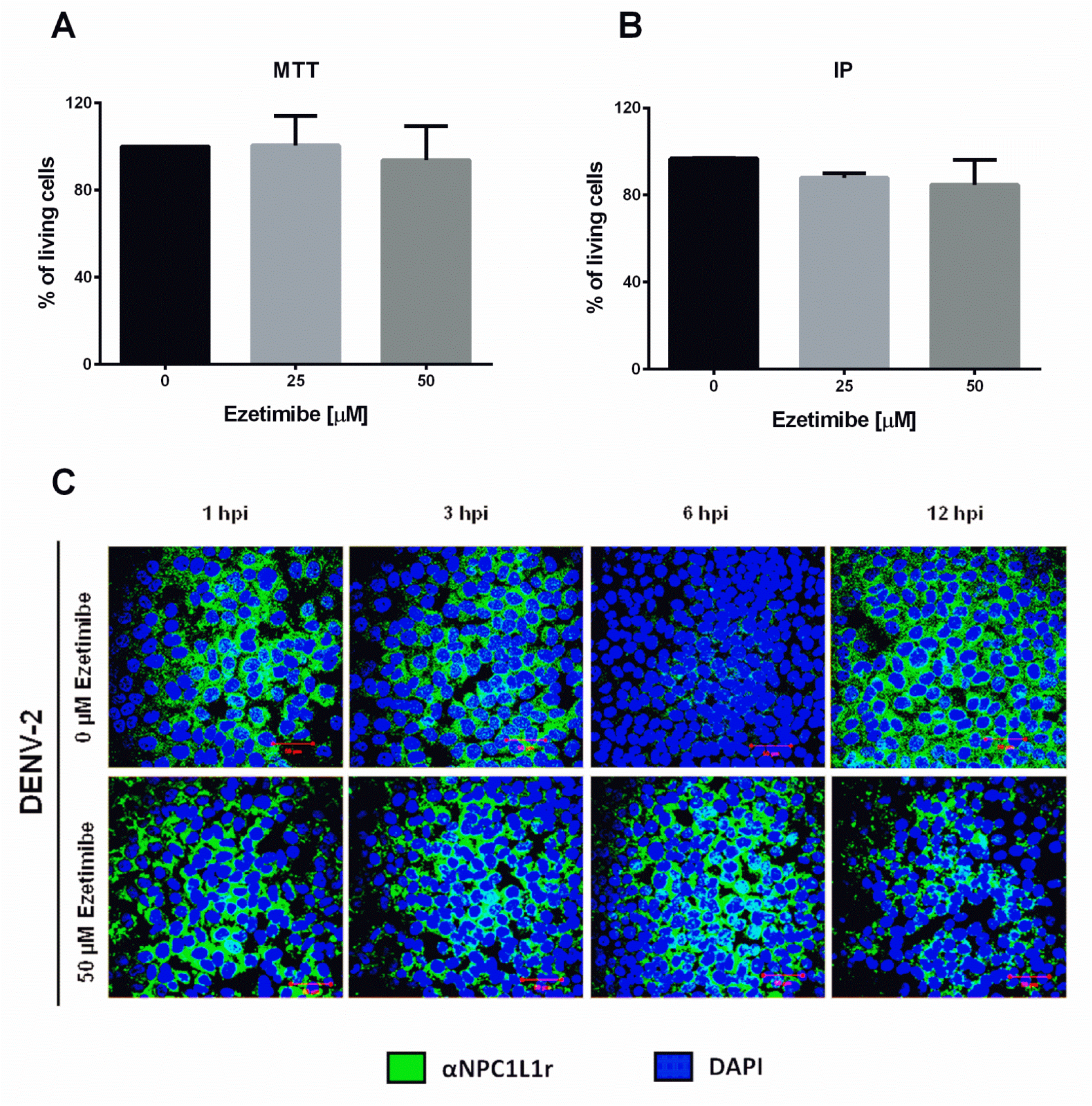
Ezetimibe does not alter cell viability and sequesters the NPC1L1 receptor on the cell membrane. Huh-7 cells were incubated in the absence of 0 (vehicle= ethanol), or in the presence (25 or 50 μM) of ezetimibe and cell viability was evaluated by incorporation of (A) propidium iodide (IP) or (B) by MTT assay. The graphs show the percentage of living cells treated for 48 hours. (C) Huh-7 cells were infected with DENV 2 and treated with 50 μM ezetimibe, fixed at 1, 3, 6 and 12 hours post-infection (hpi) and the presence of NPC1L1 receptor (green) on the cell surface was analyzed by confocal microscopy, nuclei were stained with DAPI (Blue).

**Figure 3.**
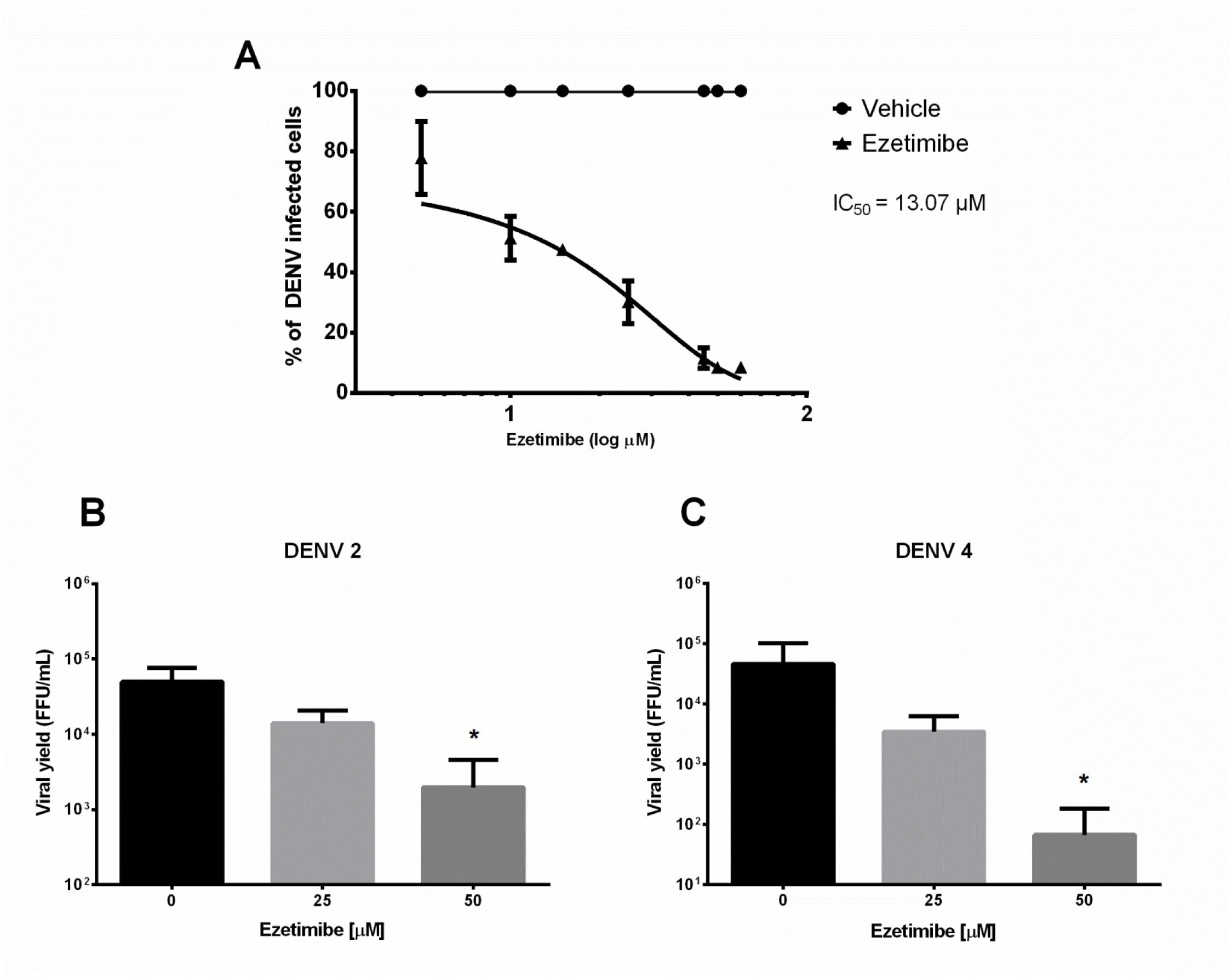
Ezetimibe has antiviral effect and reduces viral yield. (A) The IC50 was calculated over a wider range of concentrations of ezetimibe by flow cytometry. Huh7 cells infected with DENV 2 (B) or DENV 4 (C) were treated with 25 and 50 μM of ezetimibe or vehicle (0 μM) for 48 hrs. The viral titer was determined by foci forming assays. The graphs represent the percentage of infected cells as the mean and SD. The results were expressed in the log of FFU/mL from three independent assays in duplicate. \**p*= <0.05.

Remarkably, when 25 and 50 μM of ezetimibe were used, a significant reduction in viral yield, mainly at the concentration of 50 μM for DENV 2 (more than one log of UFF/mL, 95% decrease) (Fig. 3) and DENV 4 (2 logs of UFF/mL, 99% decrease) (Fig. 3 C) (p = 0.0146 and 0.0141, respectively) was detected. Since the best inhibitory effect of ezetimibe was observed at the concentration of 50 μM, this concentration was used for subsequent assays.

The next step was to evaluate the effect of ezetimibe in the number of infected cells. A significant reduction in the amount of cells labeled with the anti-prM-E antibody was observed in ezetimibe treated DENV 2 and DENV 4 infected cells compared with vehicle-treated cells by confocal microscopy (Fig. 4 A). When these differences were quantified by flow cytometry, the significant reduction in the percentage of infected cells was confirmed in ezetimibe treated cells infected with DENV2 (38.97% (*p*= 0.0140)) and DENV4 (39.43% (*p*= 0.0355)) (Fig. 4 B), compared with vehicle-treated cells. All these results confirm that the NPC1L1 receptor plays an essential role during DENV infection.

**Figure 4.**
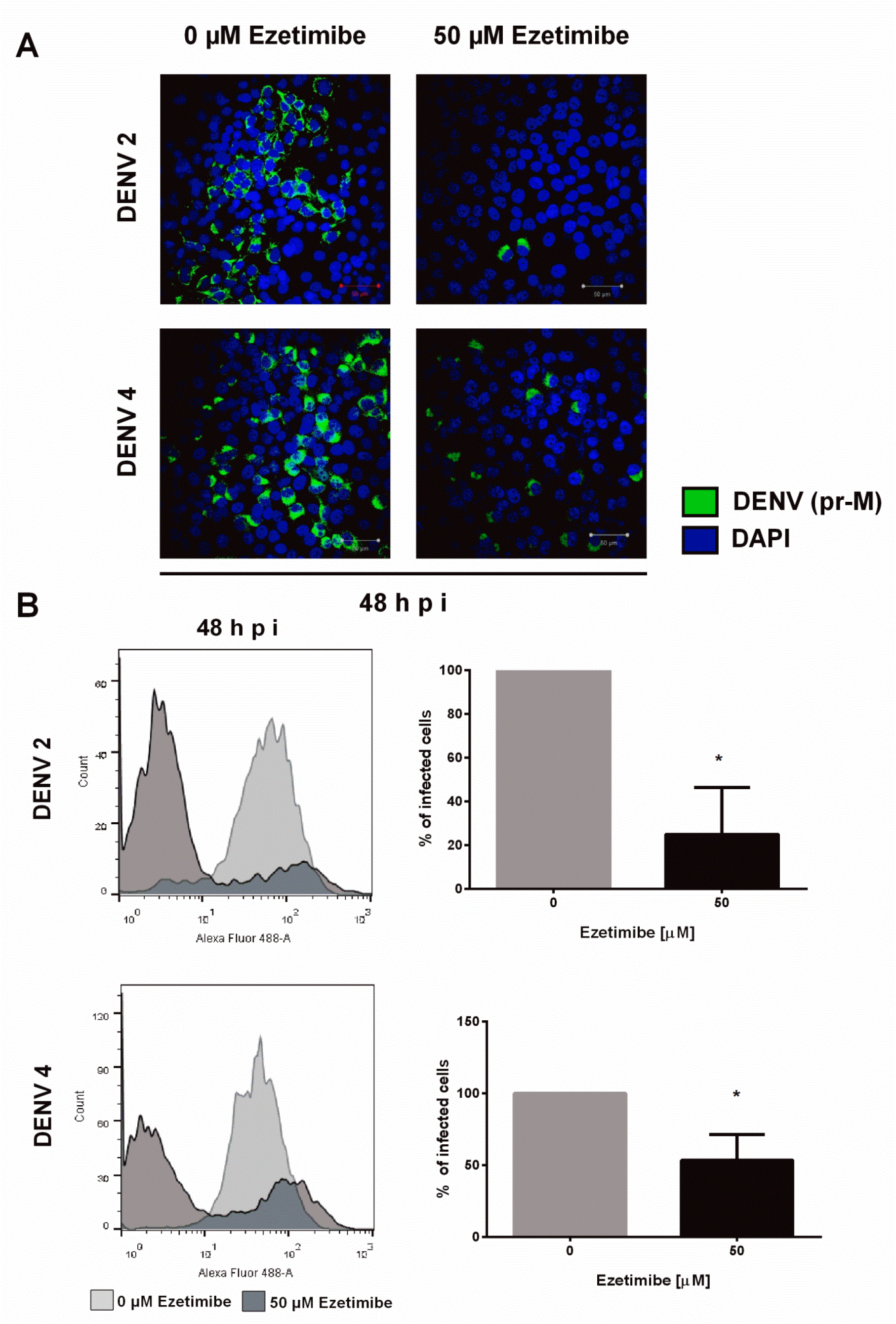
Ezetimibe induces an antiviral effect in cells infected with DENV. (A) Huh-7 cells infected with DENV 2 or DENV 4 and untreated or treated with 50 μM of ezetimibe were visualized at 48 hpi by confocal microscopy using an anti-prM-E antibody (green). Representative images of each condition are shown. (B) The percentage of cells infected with DENV 2 or 4 after treatment with 50 μM of ezetimibe was determined by flow cytometry. The results were expressed as % of infected cells of three independent assays in duplicate. \**p*= <0.05.

To exclude the possibility that the ezetimibe could have an off-target effect the effect of this drug was tested in Vero cells, which are permissive to DENV infection but do not express the NPC1L1 receptor. As it can be observed, ezetimibe did not induce a reduction in the percentage of infected cells supporting the idea that the inhibition of DENV infection induced by ezetimibe in Huh7 cells is due to the blockage of the NPC1L1 receptor (Supplemental material, Fig. 1). Moreover, the inhibition in DENV infection induced by ezetimibe in the presence of fetal calf serum (FCS), which contains cholesterol as low density (LDL) and high density (HDL) lipoproteins as well as free cholesterol (Forte et al., 1981), was less pronounced than in cells incubated in the absence of FCS, supporting the idea that the cholesterol levels in infected cells are maintained by cholesterol synthesis and also by cholesterol uptake trough at least, two receptors: LDLr and NPC1L1, because this last one was inhibited by ezetimibe (Supplemental material, Fig. 2).

### Ezetimibe inhibits post-entry steps during DENV replicative cycle

In order to investigate the specific step in DENV replicative cycle in which the NPC1L1 receptor is playing a role, the drug was added at different times of infection: 1) 6 hrs before infection (pretreatment), 2) 0, 3) 12, 4) 24 or 5) 36 hpi, according to the following scheme (Fig. 5 A). While no differences in the percentage of infected cells were observed when cells were pretreated with ezetimibe, a significant reduction in the number of infected cells was observed in cells treated with ezetimibe after infection (Fig. 5 B), supporting the idea that NPC1L1 is involved in steps after viral attachment. To analyze in further detail the specific step in which ezetimibe is inhibiting DENV infection, first, a viral binding assay in the presence of ezetimibe was performed. As shown in Figure 6 A and 6 B, the amount of virus bound to the surface of cells infected with DENV2 or DENV4 was the same in the absence or the presence of ezetimibe, suggesting that the NPC1L1 receptor not is involved in viral attachment. To confirm this result and knowing that the binding site of ezetimibe to NPC1L1 is distinct from the anti-NPC1L1 antibody recognition site (Garcia-Calvo et al., 2005; Sainz et al., 2012), the cells were incubated with the anti-NPC1L1 (Fig. 6 A and B) antibody and subsequently a viral binding assay was performed at 4 °C. DENV2 or DENV4 attachment to the cell surface was not altered by the presence of the anti-NPC1L1 antibody confirming that this receptor is not involved in viral attachment to the cell surface.

**Figure 5.**
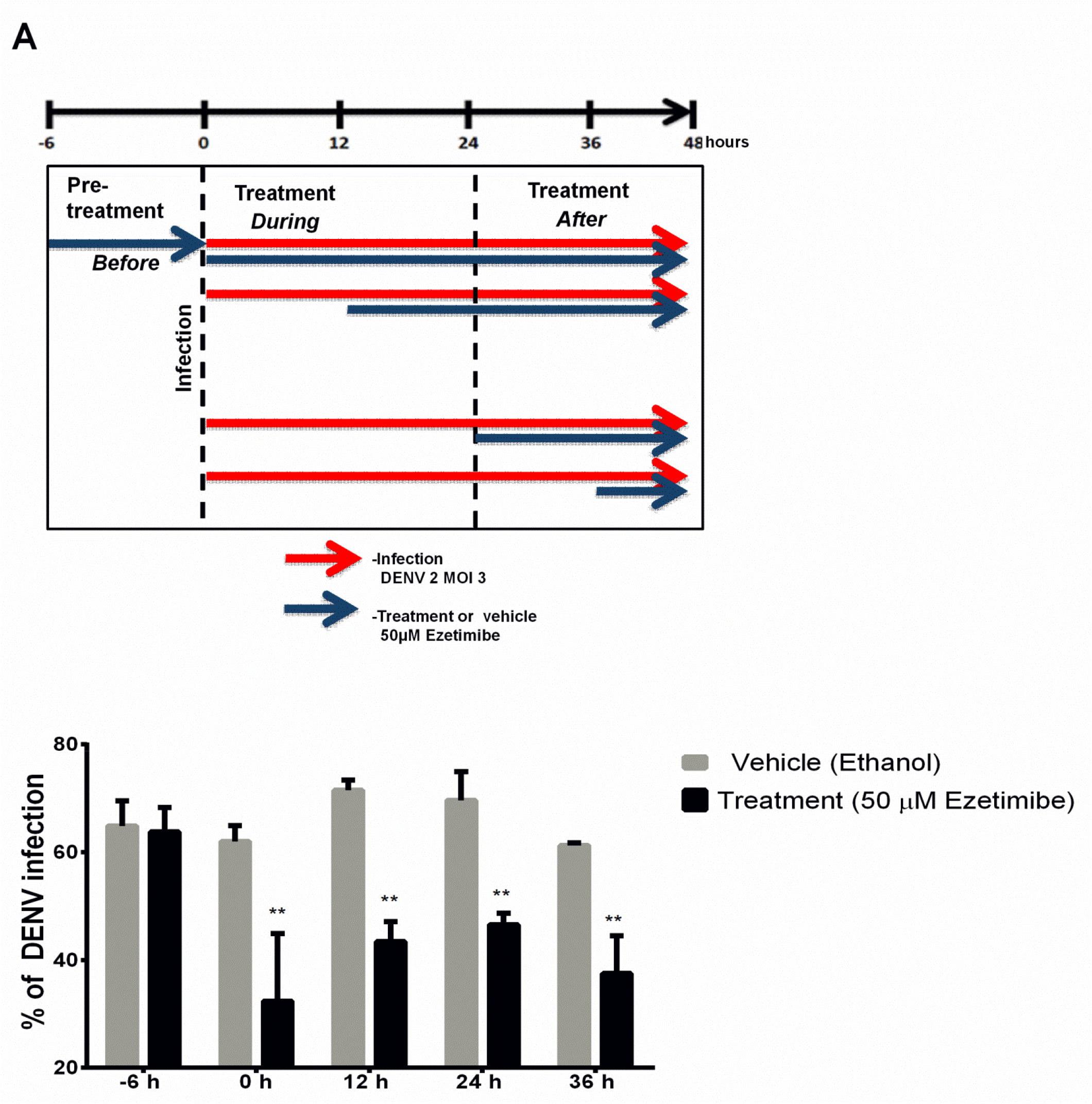
Ezetimibe inhibits DENV infection after binding and entry into Huh-7 cells. Huh-7 cells were pretreated with 50 μM for 6 hours prior to adding virus inoculum, immediately upon infection initiation (0 h), at 12 hpi, 24 hpi, and 36 hpi, represented by the scheme (A). (B) The percentage of DENV infected cells was determined by flow cytometry using the anti-prM-E antibody. The results of three independent experiments are presented. \**p*= <0.05.

**Figure 6.**
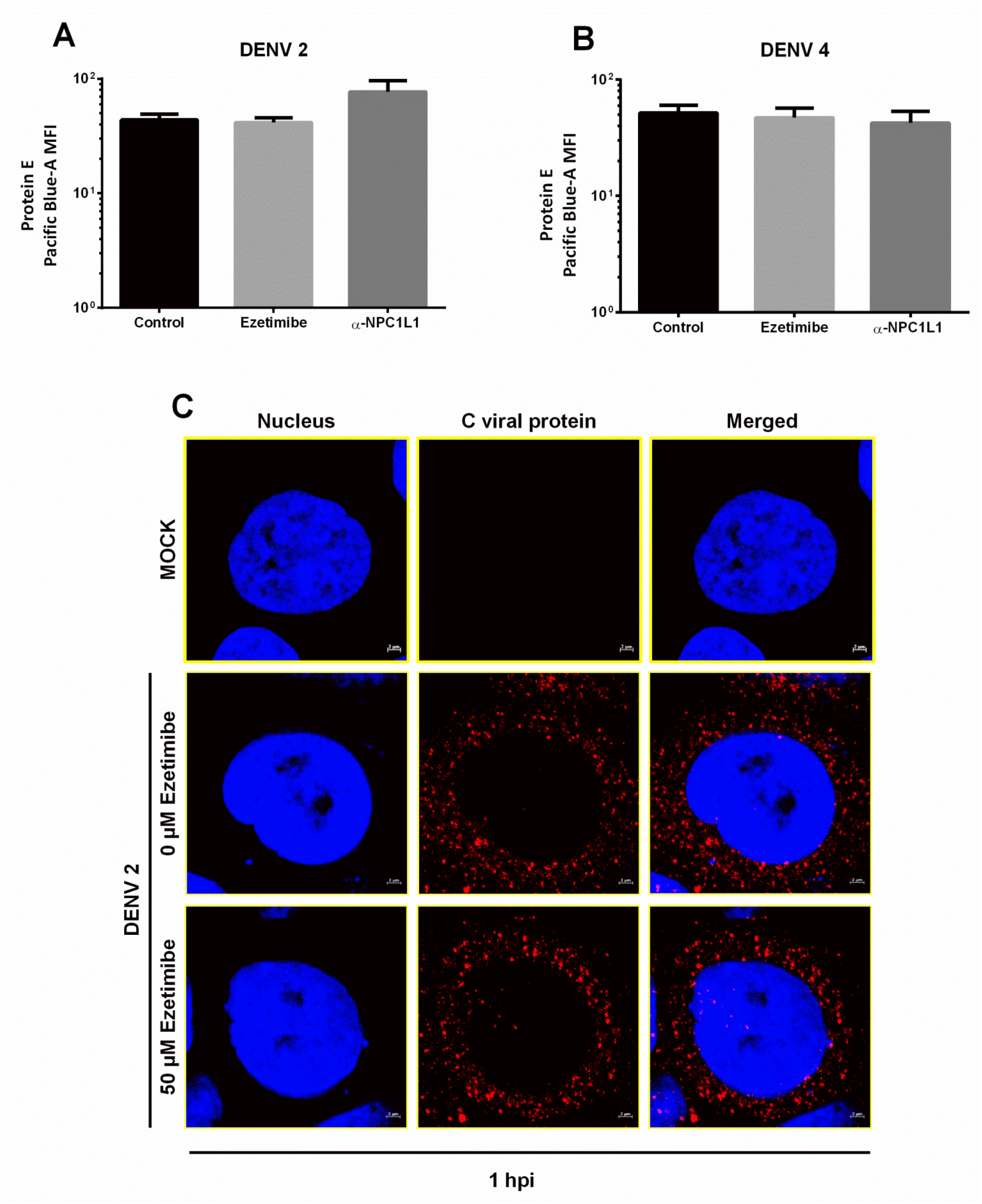
NPC1L1 is not involved in DENV attachment or entry to Huh-7 cells. In the binding assay (A and B), the Huh 7 cells in suspension were treated with ezetimibe or the anti-NPC1L1 antibody for 2 hours prior the addition of the viral inoculum. The cells were allowed to interact with DENV 2 or DENV 4 for 30 min at 4 °C, fixed and labeled with the anti-E antibody. The amount of virus bound to the surface of the cell was quantified by flow cytometry. (C) Internalization step of viral infection assay, Huh-7 cells were infected with DENV 2 and treated with 50 μM ezetimibe for two hours and fixed. The viral C protein (Red) was evaluated by confocal microscopy; nuclei were stained with DAPI (Blue). Representative imagines of each condition are shown.

To explore the possibility that ezetimibe could be disturbing the internalization of DENV, viral entry was analyzed by confocal microscopy using antibodies against C protein at a 1 hpi. The amount of C protein in the cytoplasm of infected cells was similar in ezetimibe treated and untreated cells, suggesting that the drug not alter DENV entry (Fig. 6 C).

Finally, to analyze if viral translation and/or replication could be altered by ezetimibe, the amount of viral protein and viral genome was quantified by Western blot and qRT-PCR respectively in cells untreated or treated with ezetimibe. In agreement with the results of viral yield, a significant reduction in the RNA copy number (Fig. 7 A and B) for DENV 2 and DENV 4 (p = 0.0033 and 0.0347, respectively) was detected. In the same way, a significant reduction in the amount NS3 protein (Fig. 7 C-E) from DENV 2 and DENV 4 infected cells (p = <0.0001) at both concentrations of the drug was observed, supporting the idea that ezetimibe blocks DENV infection inhibiting viral translation and replication.

**Figure 7.**
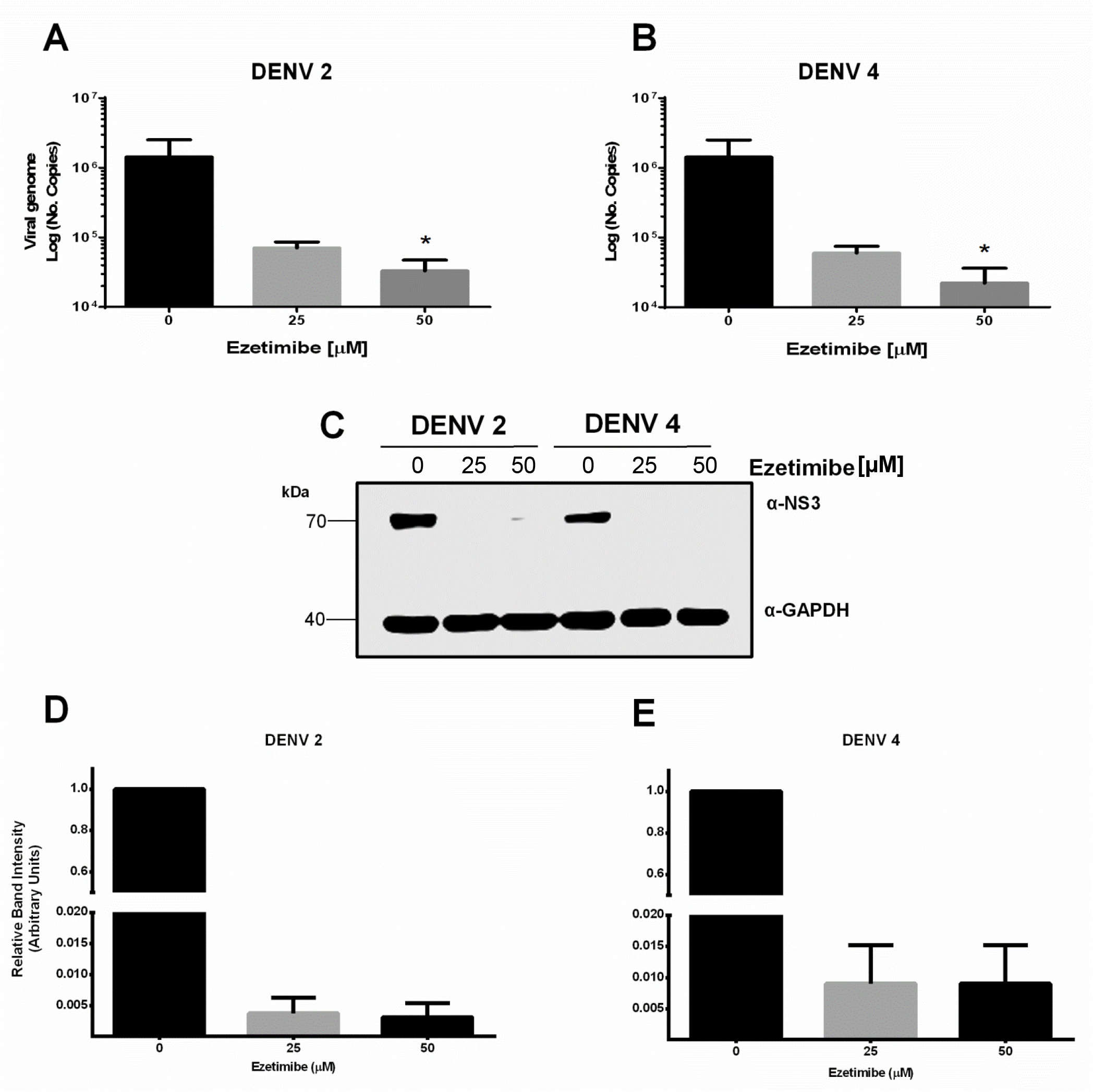
Ezetimibe reduces the number of viral RNA copies and amount of viral protein. Huh7 cells infected with DENV 2 (A, C and D) or DENV 4 (B, C, and E) were treated with 25 and 50 μM of ezetimibe or vehicle (0 μM) for 48 hrs. The viral RNA and viral NS3 protein levels we determined by qRT-PCR (A and B) and Western blot assays (C-E), respectively. The log of the number of copies of RNA ± SD and relative band intensity from three independent assays in duplicate is presented. \**p*= <0.05, \*\**p*= <0.001.

### Ezetimibe treatment affects the replicative complexes integrity

Since the main function of the NPC1L1 receptor is the uptake of cholesterol which is required for replication complexes (RC) formation, and translation and replication take place in these structures, the integrity of the RC in Huh-7 cells infected with DENV 2 and treated with ezetimibe was evaluated. The colocalization of E and NS3 proteins (Anwar et al., 2011) was analyzed by confocal microscopy (Fig. 8 A). In the absence of ezetimibe, almost 100% of the cells were infected, and a compact and perinuclear distribution of both viral proteins was observed (Fig. 8 A and B) with a positive colocalization coefficient (0.4) (Fig. 8 C). However, a substantial reduction in the amount of infected cells, as well as a diffuse perinuclear distribution of viral proteins, was observed in infected cells treated with ezetimibe (50 μM) (Fig. 8 B), with a negative colocalization coefficient (−0.02) (Fig. 8 C). These results suggest that blocking of cholesterol uptake by ezetimibe through the NPC1L1 receptor altered RC structure. To confirm the effect of ezetimibe on RC integrity, Huh-7 cells were mock infected or infected with DENV and untreated and treated for 48 h with ezetimibe and analyzed by transmission electron microscopy (TEM). The DENV-infected or vehicle-treated (0 μM ezetimibe) cells; showed membrane invaginations in the endoplasmic reticulum (ER) such as the formation of membrane packets (Vp) that contained double-membrane vesicles (Ve), and tubular structures (T), which were not found in mock-infected cells (Fig. 9 A-D). Moreover, in these infected cells virus-like particles (Vi) around membrane vesicles were observed. Vi had electron-dense nature and its size and presence of a membranous layer, strongly suggesting that it represents viral particles. Additionally, it was remarkable the observation of structures similar to ER-derived organelles called lipid droplets (LDs) (Roingeard et al., 2008) at the periphery of the viral RC (Fig. 9 D). However, in the ezetimibe DENV-infected cells with, nor the RC neither the Vi were observed. Furthermore, the ER had no alterations in its membrane, and lipid droplets were absent (Fig. 9 E and F), confirming the deleterious effect of ezetimibe in RC formation.

**Figure 8.**
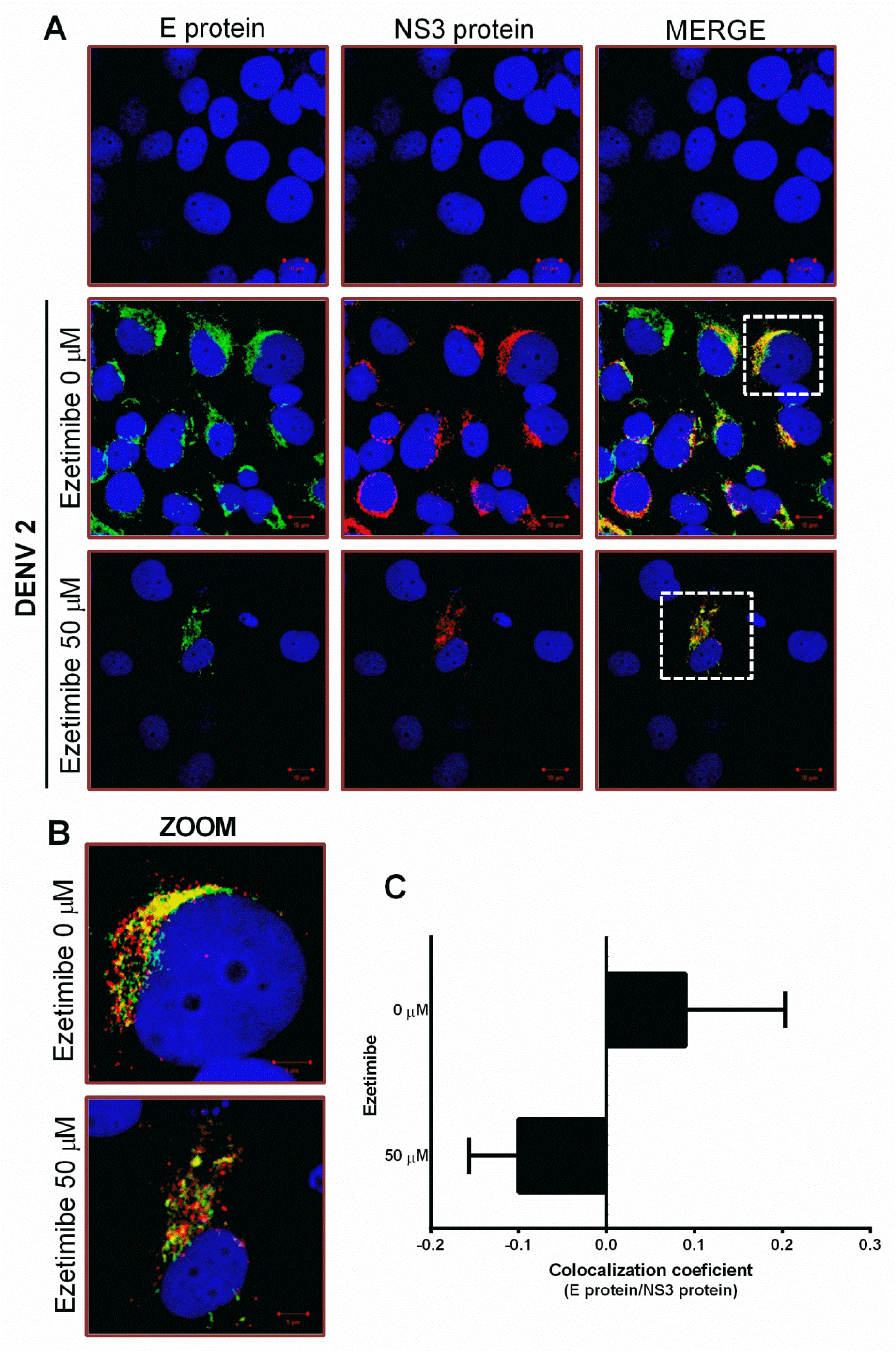
The integrity of the replicative complexes (CR) is altered in cells treated with ezetimibe. (A) The distribution of the viral proteins E (green) and NS3 (red) in DENV 2-infected Huh 7 cells and treated with vehicle (0 μM ezetimibe) or 50 μM ezetimibe was evaluated by confocal microscopy using anti-E and anti-NS3 antibodies. (B) Selected area (Dotted white box) amplified 1. 3X and 2. 2.5X showing the localization of viral proteins. The images are representatives of three independent experiments. (C) The graph compares the colocalization coefficient ±SD, through Pearson’s correlation coefficient, among both viral proteins (E protein/NS3 protein) of infected cells untreated (0 μM ezetimibe) and treated (50 μM ezetimibe).

**Figure 9.**
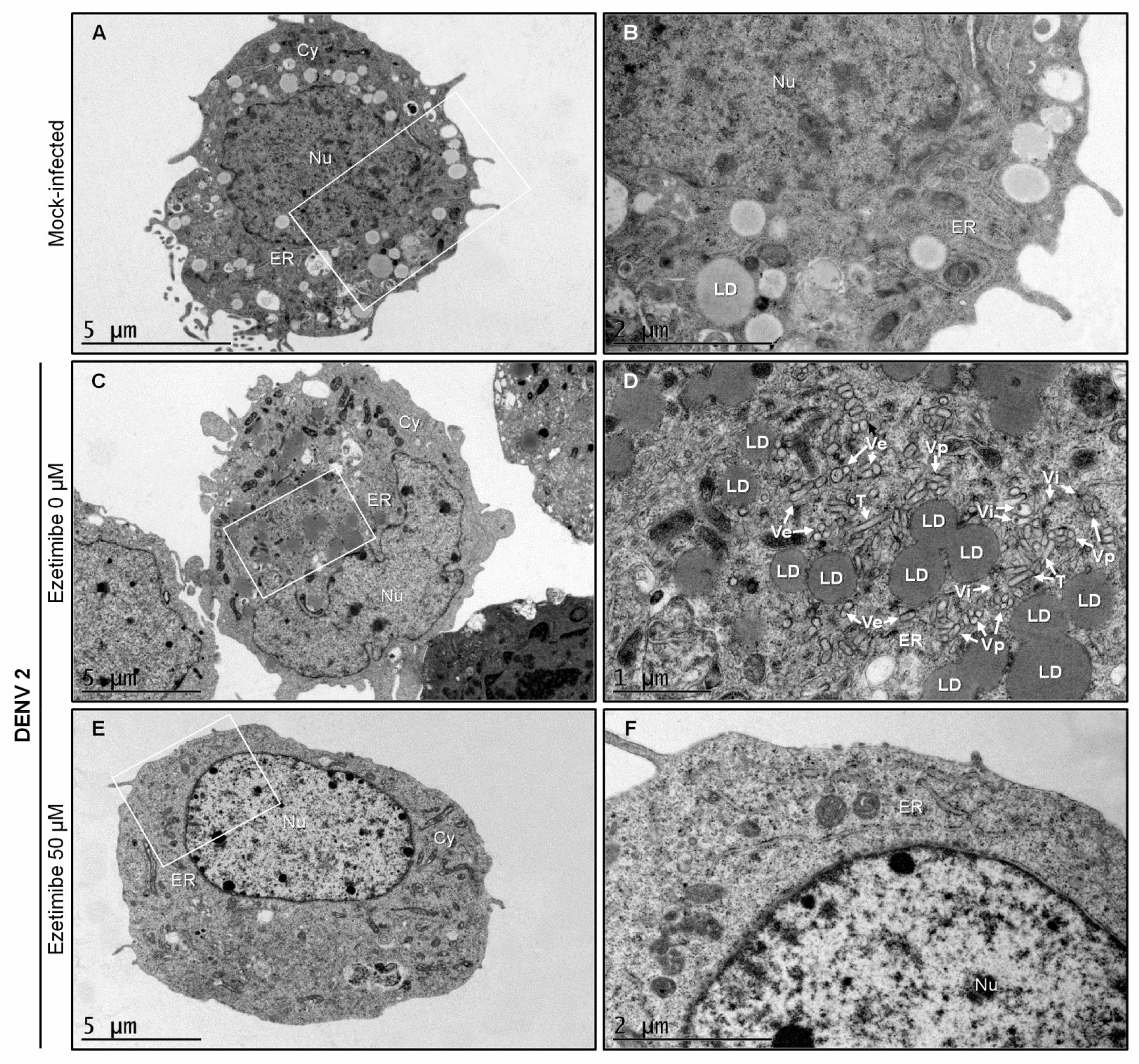
Ezetimibe affects the viral replication complex integrity. Transmission electron microscopy (TEM) analysis from Mock-infected (A and B), DENV-infected (C and D), and ezetimibe treated DENV-infected Huh-7 cells (50 μM) (E and F) at 48 hours post-infection was performed. Nu, nucleus; Cy, cytoplasm; ER, endoplasmic reticulum; Ve, double-membrane vesicles; Vp, membrane packets; T, tubular structures; Vi, virus-like particles; and LD, lipid droplet. In the TEM analysis were assessed 100 cells profiles for each experimental condition.

This alteration in the RC formation is likely responsible for the reduction in the translation and replication inhibition observed in the ezetimibe infected and treated cells. To confirm that the blockage of NPC1L1 receptor causes a reduction in cholesterol accumulation in the infected cells, the levels of cholesterol present in DENV-infected cells were evaluated in untreated, and ezetimibe treated cells for 48 h by confocal microscopy and by a fluorometric assay. As it was described previously (Soto-Acosta et al., 2013), an increase in cholesterol levels was observed in infected cells compared to MOCK-infected cells (Fig. 10 A-C). Moreover, filipin III dyed cholesterol was detected collocating with the non-structural protein 4A (NS4A) (a component of the RC (Miller et al., 2007)), in the perinuclear region. However, in the ezetimibe treated infected cells, a reduction in the amount of cholesterol (Fig. 10A-C) as well as in the number of infected cells was observed.

**Figure 10.**
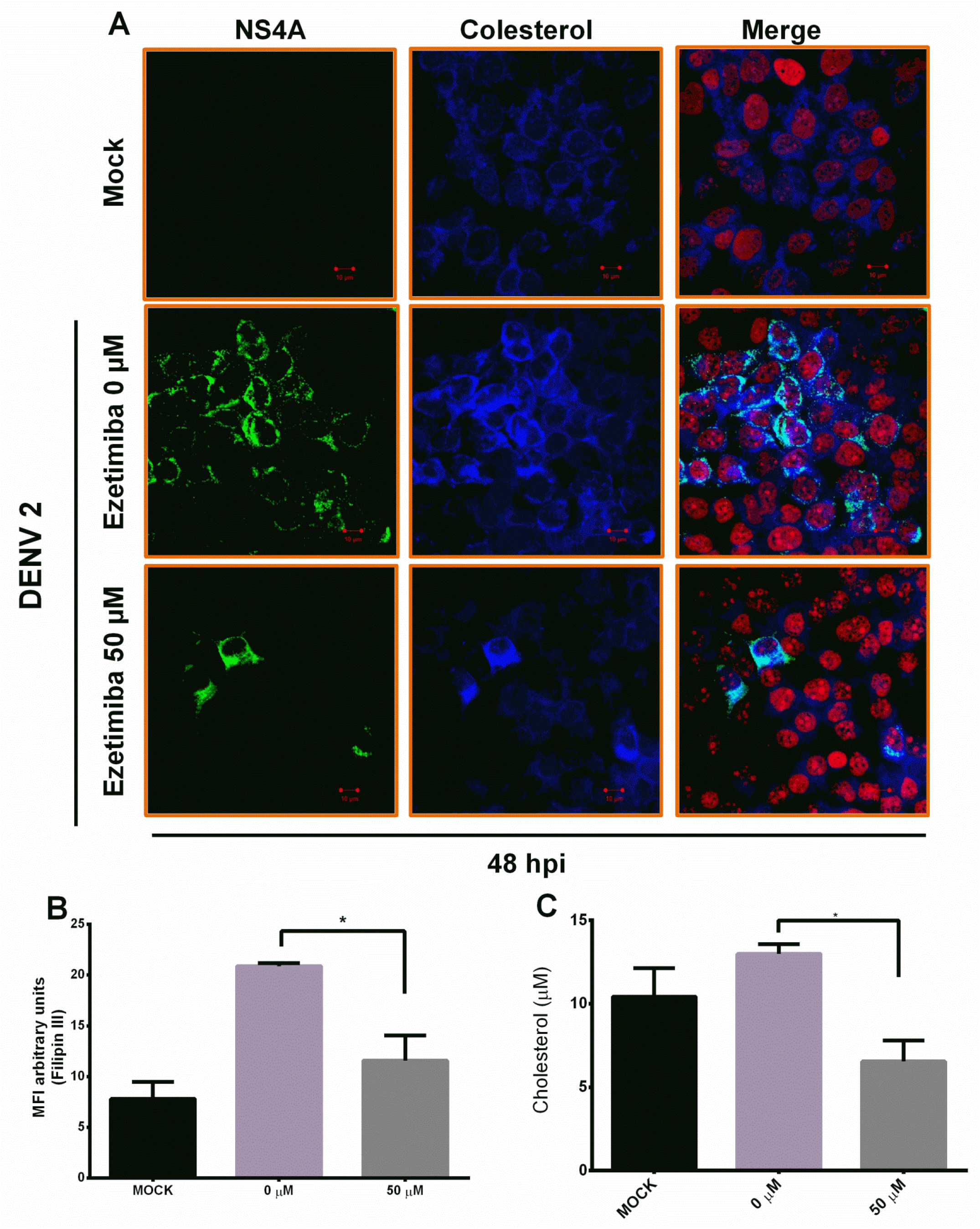
Ezetimibe reduces cellular cholesterol and disrupts the replication complexes formation. Uninfected (MOCK) and infected (48 h) Huh-7 cells untreated (0 μM) or treated with ezetimibe (50μM) were incubated with the Filipin III complex (blue) and anti-NS4A antibody (green) and were analyzed by confocal microscopy. The nuclei were stained with propidium iodide (red). (B) Cholesterol levels were evaluated by pixels analysis, showing the mean fluorescence intensity (MFI) of Filipin III complex, and by a (C) fluorometric enzymatic assay. \**p*= <0.05, \*\**p*= <0.01.

All these results indicate that NPC1L1 receptor is involved in cholesterol accumulation required for RC formation during DENV infection. Thus, ezetimibe represents an effective antiviral drug for DENV infection.

## Discussion

The opportunity offered by host-directed antiviral therapy (HDA), targeting host factors that are usurped by DENV for replication, is promising. Given the genetic austerity of the viruses, it depends on cellular factors and organelles to complete its viral cycle. One of the cellular components required during DENV infection is the cholesterol (Faustino et al., 2014; Soto-Acosta et al., 2013; Villareal et al., 2015). Moreover, changes in serum lipid profile have been correlated with clinical manifestations in DENV infected patients (Biswas et al., 2015; Durán et al., 2015; Osuna-Ramos et al., 2018; Suvarna and Rane, 2009; van Gorp et al., 2002). Thus, there is sufficient evidence to conclude that cholesterol and lipoproteins have a fundamental role during DENV infection (Faustino et al., 2014; Soto-Acosta et al., 2013; Villareal et al., 2015). Cholesterol is an important component of the cell membranes and is essential for the adequate cellular functioning (Fernández et al., 2004; Simons and Ikonen, 2000). Cholesterol levels in the cells are controlled by biosynthesis, efflux from cells, and uptake (Simons and Ikonen, 2000). Previous studies carried out in our laboratory have shown that the increase in cholesterol levels in infected cells are related to 1) an increased activity of HMG-CoA (3-hydroxy-3-methylglutaryl-coenzyme A) reductase enzyme, through the dephosphorylation of AMPK (AMP-activated protein kinase); and with 2) an augment in cholesterol uptake by the increase in the amount of LDLr on the cell surface, (Soto-Acosta et al., 2017, 2013). Therefore, treatment with statins, which inhibit the activity of HMG-CoA reductase, or metformin which activates APMK in cellular and/or animal models appears promising against DENV infection (Martinez-Gutierrez et al., 2014; Rothwell et al., 2009b; Soto-Acosta et al., 2017, 2013). However, while lovastatin treatment was not able to inhibit DENV infection in a clinical trial in humans (Whitehorn et al., 2016), the metformin reduced clinical complications caused by DENV infection in diabetic patients (Htun et al., 2018), supporting the idea that the cholesterol is required during DENV infection. Since cholesterol is not only obtained by an increase in cholesterol synthesis but also through cellular receptors such as the LDLr (Soto-Acosta et al., 2013) the function of other cholesterol receptors such as NPC1L1 receptor in DENV infection was evaluated. Interestingly, in our study, we observed an increase of the NPC1L1 receptor on the cell surface during the first hour after infection as it was described previously with the LDLr (Soto-Acosta et al., 2013). Subsequently, at 3 and 6 hours post-infection a reduction of the NPC1L1 receptor on the cell surface was observed (Fig. 1 A and B). However, unlike what was reported with LDLr in DENV infection, at 12 hpi we observed recovery of the NPC1L1 receptor on the cell membrane. At this time, there is a significant viral replication activity with important cholesterol requirements. It has been described that when cholesterol is removed from the medium and later added to liver cells, 85% of the NPC1L1 receptor is located in the plasma membrane to capture it (Ge et al., 2008). This was observed in the DENV-infected cells when the cells require a higher amount of cholesterol, and it is uptake from the media at early times. This finding suggests that binding or entry of DENV triggers cholesterol accumulation in the cell via LDLr and NPC1L1 receptor, but during viral replication, the cholesterol requirement increases and it is obtained through the NPC1L1 receptor uptake and by the increase in cholesterol synthesis by the HMG-CoA reductase (Soto-Acosta et al., 2017). It is known that the NPC1L1 receptor is involved in the uptake of cholesterol in liver cells, which can be blocked by ezetimibe, a drug approved by the FDA as cholesterol-lowering therapy (Garcia-Calvo et al., 2005). Our results suggest that ezetimibe can reduce the production of new viral particles (Fig. 3 B and C), inhibits the DENV genome synthesis (Fig. 7 A and B) and reduce viral protein synthesis (Fig. 7 C-E) of DENV 2 and DENV 4 by inhibiting the uptake of extracellular cholesterol and blocking the NPC1L1 receptor by arresting it in the cell membrane during the first 12 hours of infection (Fig. 2 C). In addition, we were able to exclude that ezetimibe could be having an off-target effect since it was unable to inhibit DENV infection in Vero cells which lack the NPC1L1 receptor.

Since treatment with ezetimibe was not able to inhibit DENV binding or entry, we can suggest that the NPC1L1 receptor is not an entry factor for DENV. This was also the case for hepatitis B virus (HBV), in which ezetimibe blocked post-entry events but not binding or entry (Lucifora et al., 2013). In contrast, ezetimibe inhibited attachment and entry of Hepatitis C virus (HCV) considering to the NPC1L1 receptor as an entry factor (Sainz et al., 2012). Interestingly, ezetimibe had a higher potential to inhibit DENV than HCV and HBV infections (Lucifora et al., 2013; Sainz et al., 2012), because the in vitro IC50 for DENV (13.07 μM) was lower than the one reported for HBV and HCV, supporting the idea that ezetimibe is a good candidate as an anti-DENV drug. It is important to notice that the reduction in viral yield, the inhibition in the amount of viral genome and proteins after treatment with ezetimibe, can be considered as a consequence of the loss of the integrity of RC (Fig. 8 and 9), because a reduction in cholesterol levels, causes an inhibition in RC formation in DENV-infected cells (Anwar et al., 2011; Miller et al., 2007). The confocal microscopy and TEM analysis demonstrated that ezetimibe affects the integrity of the membrane vesicles reported as RC in DENV infected cells (Junjhon et al., 2014; Reyes-Ruiz et al., 2018; Welsch et al., 2009). Additionally, a large number of LDs at the periphery of the viral RC observed in infected cells (Samsa et al., 2009) were also disrupted in the presence of ezetimibe. Thus, the life cycle of LDs, which starts when fatty acids and sterols enter into the cell (Guo et al., 2009; Rachid et al., 2013), is blocked by Ezetimibe reducing the LDs formation. A disruption of RC has also been observed in DENV-infected cells treated with drugs with hypolipidemic properties such as NDGA which causes a reduction in the colocalization of the viral proteins E and NS3 in the perinuclear region (Soto-Acosta et al., 2014). On the other hand, since DENV infection stimulates the accumulation of cholesterol in RC through the activation of HMG-CoA reductase, drugs such as metformin (AMPK activator that reduces HMG-CoA reductase activity) and lovastatin (HMG-CoA reductase inhibitor), both with hypolipemic activity, decreased the colocalization of the non-structural protein 4A (NS4A) with cellular cholesterol (stained with filipin III) (Soto-Acosta et al., 2017), corroborating the importance of cholesterol in the formation and integrity of RC during DENV infection. The fact that ezetimibe reduces the cellular cholesterol levels confirms the importance of exogenous cholesterol in the integrity and formation of RC during DENV replication.

In summary, in the present study, we demonstrate that the NPC1L1 receptor plays an essential role in the cholesterol accumulation during DENV infection. It is possible that this receptor is working in collaboration with the LDLr to uptake cholesterol from media early after DENV infection. Since there are multiple other cellular receptors responsible for cholesterol uptake, such as the scavenger receptor class B type I (SR-BI) receptor (Li et al., 2013), the participation of those receptors need to be evaluated.

Thus, our results indicate that the antiviral effect induced by ezetimibe is caused by a reduction of cholesterol uptake, which avoid the RC formation reducing the viral genome replication and translation and the formation of new viral particles. These results together with our previous findings allow us to conclude that during DENV infection there is an increase in cholesterol levels due to an increase in cholesterol synthesis and an increase in cholesterol uptake through two different receptors LDL (Soto-Acosta et al., 2013) and NPC1L1 receptors. Finally, the requirement of extracellular cholesterol during DENV infection can be exploited with the use of cholesterol-lowering drugs. Since ezetimibe is a FDA-approved drug, it can be a suitable candidate for an HDA therapy for DENV infection.

## Acknowledgments

The authors thank to Dr. Bibiana Chávez-Munguía and Anel E. Lagunes Guillen for their valuable help, assistance, and preparation of the electron microscopy image samples and to Fernando Medina and Jaime Zarco for their technical assistance. This work was supported by CONACYT (Mexico) grant 220824, and Fundación Miguel Alemán. The funders had no role in study design, data collection, and analysis, decision to publish, or preparation of the manuscript.

---

**Supplemental material, Figure 1.**
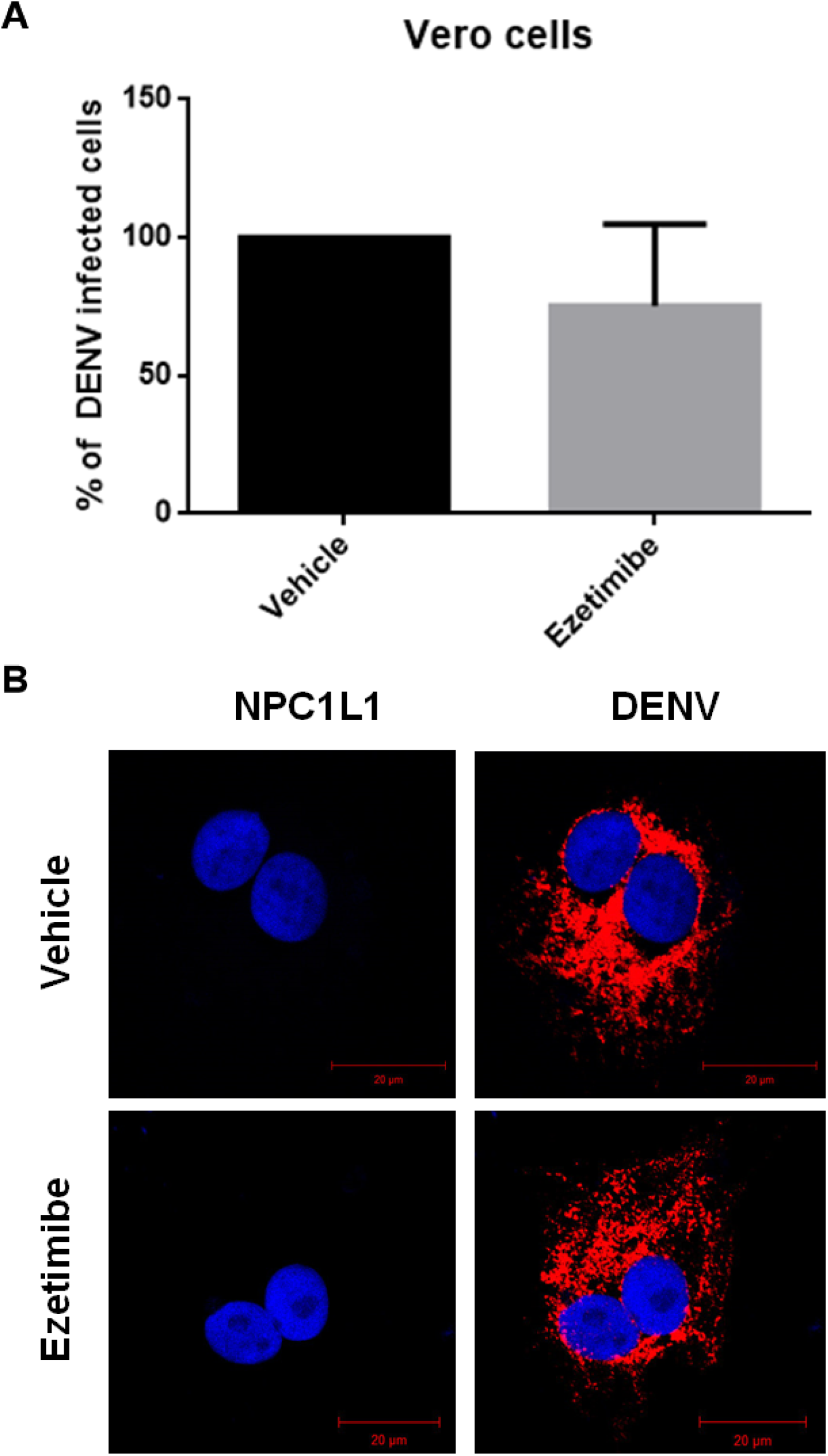
Off-target effects of ezetimibe in DENV infection. Vero cells were infected with DENV 2 at MOI 3 and treated with 50 μM ezetimibe. The cells were fixed at 48 hours post-infection, and the percentage of infected cells was analyzed by (A) flow cytometry and (B) confocal microscopy by staining the cells with NS4 viral protein (red) and NPC1L1.

**Supplemental material, Figure 2.**
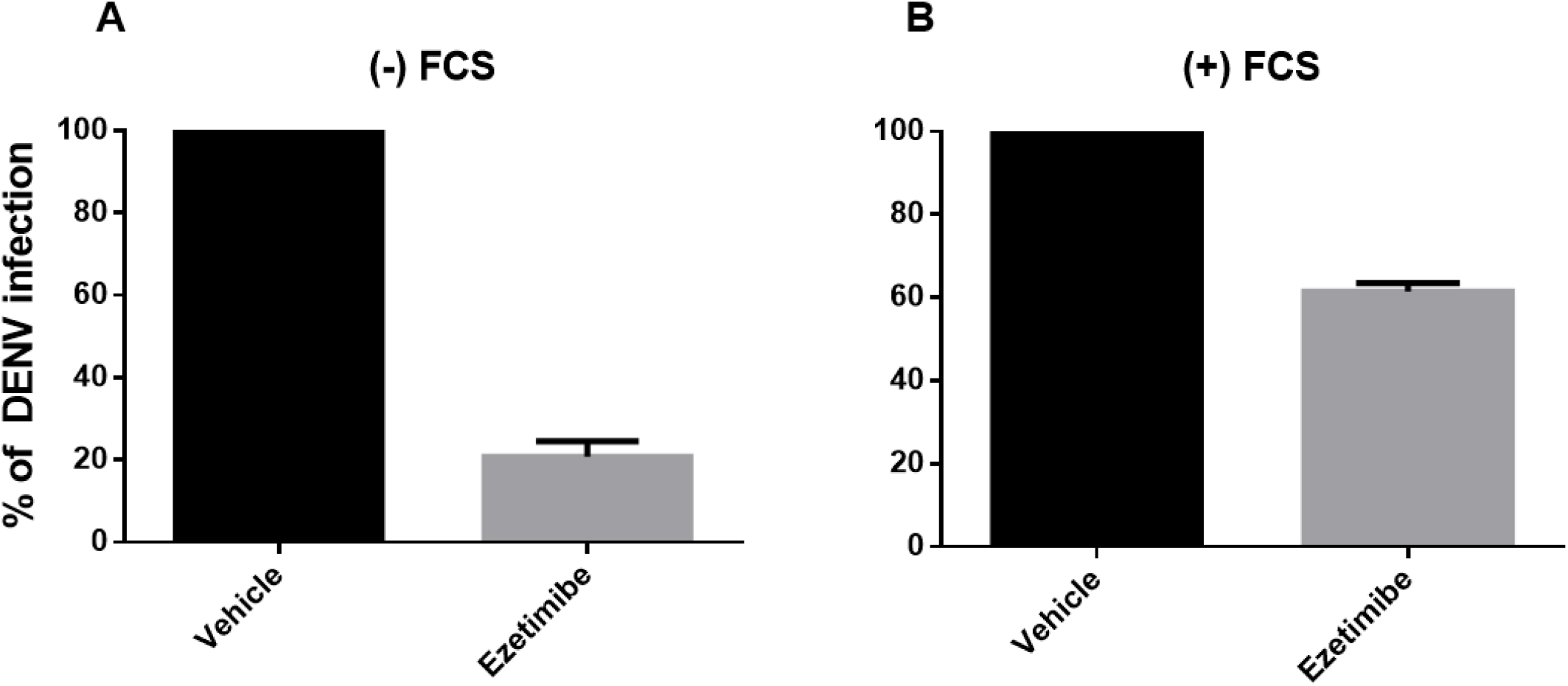
Fetal calf serum (FCS)-favors DENV infection. Huh-7 cells were infected with DENV 2 at MOI 3, then the cells were treated with 50 μM of ezetimibe diluted in FCS-free (A) or non-FCS free (B) medium for 12 h, the cells were fixed and the number of infected cells was evaluated by flow cytometry. The percentage of infection was normalized with the vehicle.

## References

Acosta, E.G., Bartenschlager, R., 2016. The quest for host targets to combat dengue virus infections. Curr. Opin. Virol. 20, 47–54. https://doi.org/10.1016/j.coviro.2016.09.003

Anwar, A., Hosoya, T., Leong, K.M., Onogi, H., Okuno, Y., Hiramatsu, T., Koyama, H., Suzuki, M., Hagiwara, M., Garcia-Blanco, M.A., 2011. The Kinase Inhibitor SFV785 Dislocates Dengue Virus Envelope Protein from the Replication Complex and Blocks Virus Assembly. PLoS ONE 6. https://doi.org/10.1371/journal.pone.0023246

Bautista-Carbajal, P., Soto-Acosta, R., Angel-Ambrocio, A.H., Cervantes-Salazar, M., Loranca-Vega, C.I., Herrera-Martínez, M., Del Angel, R.M., 2017. The calmodulin antagonist W-7 (N-(6-aminohexyl)-5-chloro-1-naphthalenesulfonamide hydrochloride) inhibits DENV infection in Huh-7 cells. Virology 501, 188–198. https://doi.org/10.1016/j.virol.2016.12.004

Betters, J.L., Yu, L., 2010. NPC1L1 and cholesterol transport. FEBS Lett. 584, 2740–2747. https://doi.org/10.1016/j.febslet.2010.03.030

Biswas, H.H., Gordon, A., Nuñez, A., Perez, M.A., Balmaseda, A., Harris, E., 2015. Lower Low-Density Lipoprotein Cholesterol Levels Are Associated with Severe Dengue Outcome. PLoS Negl. Trop. Dis. 9, e0003904. https://doi.org/10.1371/journal.pntd.0003904

Carro, A.C., Damonte, E.B., 2013. Requirement of cholesterol in the viral envelope for dengue virus infection. Virus Res. 174, 78–87. https://doi.org/10.1016/j.virusres.2013.03.005

Chang, T.-Y., Chang, C., 2008. Ezetimibe blocks internalization of the NPC1L1/cholesterol complex. Cell Metab. 7, 469–471. https://doi.org/10.1016/j.cmet.2008.05.001

Durán, A., Carrero, R., Parra, B., González, A., Delgado, L., Mosquera, J., Valero, N., 2015. Association of lipid profile alterations with severe forms of dengue in humans. Arch. Virol. 160, 1687–1692. https://doi.org/10.1007/s00705-015-2433-z

Faustino, A.F., Carvalho, F.A., Martins, I.C., Castanho, M.A.R.B., Mohana-Borges, R., Almeida, F.C.L., Da Poian, A.T., Santos, N.C., 2014. Dengue virus capsid protein interacts specifically with very low-density lipoproteins. Nanomedicine Nanotechnol. Biol. Med. 10, 247–255. https://doi.org/10.1016/j.nano.2013.06.004

Fernández, C., Lobo Md, M. del V.T., Gómez-Coronado, D., Lasunción, M.A., 2004. Cholesterol is essential for mitosis progression and its deficiency induces polyploid cell formation. Exp. Cell Res. 300, 109–120. https://doi.org/10.1016/j.yexcr.2004.06.029

Forte, T.M., Bell-Quint, J.J., Cheng, F., 1981. Lipoproteins of fetal and newborn calves and adult steer: A study of developmental changes. Lipids 16, 240–245. https://doi.org/10.1007/BF02535023s

Garcia-Calvo, M., Lisnock, J., Bull, H.G., Hawes, B.E., Burnett, D.A., Braun, M.P., Crona, J.H., Davis, H.R., Dean, D.C., Detmers, P.A., Graziano, M.P., Hughes, M., Macintyre, D.E., Ogawa, A., O’neill, K.A., Iyer, S.P.N., Shevell, D.E., Smith, M.M., Tang, Y.S., Makarewicz, A.M., Ujjainwalla, F., Altmann, S.W., Chapman, K.T., Thornberry, N.A., 2005. The target of ezetimibe is Niemann-Pick C1-Like 1 (NPC1L1). Proc. Natl. Acad. Sci. U. S. A. 102, 8132–8137. https://doi.org/10.1073/pnas.0500269102

Ge, L., Wang, J., Qi, W., Miao, H.-H., Cao, J., Qu, Y.-X., Li, B.-L., Song, B.-L., 2008. The cholesterol absorption inhibitor ezetimibe acts by blocking the sterol-induced internalization of NPC1L1. Cell Metab. 7, 508–519. https://doi.org/10.1016/j.cmet.2008.04.001

Guzman, M.G., Harris, E., 2015. Dengue. The Lancet 385, 453–465. https://doi.org/10.1016/S0140-6736(14)60572-9

Hadinegoro, S.R.S., 2012. The revised WHO dengue case classification: does the system need to be modified? Paediatr. Int. Child Health 32, 33–38. https://doi.org/10.1179/2046904712Z.00000000052

Halstead, S.B., Russell, P.K., 2016. Protective and immunological behavior of chimeric yellow fever dengue vaccine. Vaccine 34, 1643–1647. https://doi.org/10.1016/j.vaccine.2016.02.004

Hasan, S., Jamdar, S.F., Alalowi, M., Al Ageel Al Beaiji, S.M., 2016. Dengue virus: A global human threat: Review of literature. J. Int. Soc. Prev. Community Dent. 6, 1–6. https://doi.org/10.4103/2231-0762.175416

Htun, H.L., Yeo, T.W., Tam, C.C., Pang, J., Leo, Y.S., Lye, D.C., 2018. Metformin Use and Severe Dengue in Diabetic Adults. Sci. Rep. 8, 3344. https://doi.org/10.1038/s41598-018-21612-6

Jia, L., Betters, J.L., Yu, L., 2011. Niemann-pick C1-like 1 (NPC1L1) protein in intestinal and hepatic cholesterol transport. Annu. Rev. Physiol. 73, 239–259. https://doi.org/10.1146/annurev-physiol-012110-142233

Junjhon, J., Pennington, J.G., Edwards, T.J., Perera, R., Lanman, J., Kuhn, R.J., 2014. Ultrastructural characterization and three-dimensional architecture of replication sites in dengue virus-infected mosquito cells. J. Virol. 88, 4687–4697. https://doi.org/10.1128/JVI.00118-14

Li, Y., Kakinami, C., Li, Q., Yang, B., Li, H., 2013. Human apolipoprotein A-I is associated with dengue virus and enhances virus infection through SR-BI. PloS One 8, e70390. https://doi.org/10.1371/journal.pone.0070390

Lim, S.P., Wang, Q.-Y., Noble, C.G., Chen, Y.-L., Dong, H., Zou, B., Yokokawa, F., Nilar, S., Smith, P., Beer, D., Lescar, J., Shi, P.-Y., 2013. Ten years of dengue drug discovery: progress and prospects. Antiviral Res. 100, 500–519. https://doi.org/10.1016/j.antiviral.2013.09.013

Lucifora, J., Esser, K., Protzer, U., 2013. Ezetimibe blocks hepatitis B virus infection after virus uptake into hepatocytes. Antiviral Res. 97, 195–197. https://doi.org/10.1016/j.antiviral.2012.12.008

Martinez-Gutierrez, M., Correa-Londoño, L.A., Castellanos, J.E., Gallego-Gómez, J.C., Osorio, J.E., 2014. Lovastatin delays infection and increases survival rates in AG129 mice infected with dengue virus serotype 2. PloS One 9, e87412. https://doi.org/10.1371/journal.pone.0087412

Miller, S., Kastner, S., Krijnse-Locker, J., Bühler, S., Bartenschlager, R., 2007. The non-structural protein 4A of dengue virus is an integral membrane protein inducing membrane alterations in a 2K-regulated manner. J. Biol. Chem. 282, 8873–8882. https://doi.org/10.1074/jbc.M609919200

Osuna-Ramos, J.F., Rendón-Aguilar, H., Reyes-Ruiz, J.M., Del Ángel, R.M., Romero-Utrilla, A., Ríos-Burgueño, E.R., Velarde-Rodriguez, I., Velarde-Félix, J.S., 2018. The correlation of TNF alpha levels with the lipid profile of dengue patients. J. Med. Virol. https://doi.org/10.1002/jmv.25056

Reyes-Ruiz, J.M., Osuna-Ramos, J.F., Cervantes-Salazar, M., Lagunes Guillen, A.E., Chávez-Munguía, B., Salas-Benito, J.S., Del Ángel, R.M., 2018. Strand-like structures and the nonstructural proteins 5, 3 and 1 are present in the nucleus of mosquito cells infected with dengue virus. Virology 515, 74–80. https://doi.org/10.1016/j.virol.2017.12.014

Roingeard, P., Hourioux, C., Blanchard, E., Prensier, G., 2008. Hepatitis C virus budding at lipid droplet-associated ER membrane visualized by 3D electron microscopy. Histochem. Cell Biol. 130, 561–566. https://doi.org/10.1007/s00418-x008-0447-2

Rothwell, C., Lebreton, A., Young Ng, C., Lim, J.Y.H., Liu, W., Vasudevan, S., Labow, M., Gu, F., Gaither, L.A., 2009. Cholesterol biosynthesis modulation regulates dengue viral replication. Virology 389, 8–19. https://doi.org/10.1016/j.virol.2009.03.025

Sainz, B., Barretto, N., Martin, D.N., Hiraga, N., Imamura, M., Hussain, S., Marsh, K.A., Yu, X., Chayama, K., Alrefai, W.A., Uprichard, S.L., 2012. Identification of the Niemann-Pick C1-like 1 cholesterol absorption receptor as a new hepatitis C virus entry factor. Nat. Med. 18, 281–285. https://doi.org/10.1038/nm.2581

Samsa, M.M., Mondotte, J.A., Iglesias, N.G., Assunção-Miranda, I., Barbosa-Lima, G., Da Poian, A.T., Bozza, P.T., Gamarnik, A.V., 2009. Dengue virus capsid protein usurps lipid droplets for viral particle formation. PLoS Pathog. 5, e1000632. https://doi.org/10.1371/journal.ppat.1000632

Simmons, C.P., Farrar, J.J., Nguyen, van V.C., Wills, B., 2012. Dengue. N. Engl. J. Med. 366, 1423–1432. https://doi.org/10.1056/NEJMra1110265

Simons, K., Ikonen, E., 2000. How cells handle cholesterol. Science 290, 1721–1726.

Soto-Acosta, R., Bautista-Carbajal, P., Cervantes-Salazar, M., Angel-Ambrocio, A.H., Del Angel, R.M., 2017. DENV up-regulates the HMG-CoA reductase activity through the impairment of AMPK phosphorylation: A potential antiviral target. PLoS Pathog. 13, e1006257. https://doi.org/10.1371/journal.ppat.1006257

Soto-Acosta, R., Bautista-Carbajal, P., Syed, G.H., Siddiqui, A., Del Angel, R.M., 2014. Nordihydroguaiaretic acid (NDGA) inhibits replication and viral morphogenesis of dengue virus. Antiviral Res. 109, 132–140. https://doi.org/10.1016/j.antiviral.2014.07.002

Soto-Acosta, R., Mosso, C., Cervantes-Salazar, M., Puerta-Guardo, H., Medina, F., Favari, L., Ludert, J.E., del Angel, R.M., 2013. The increase in cholesterol levels at early stages after dengue virus infection correlates with an augment in LDL particle uptake and HMG-CoA reductase activity. Virology 442, 132–147. https://doi.org/10.1016/j.virol.2013.04.003

Suvarna, J.C., Rane, P.P., 2009. Serum lipid profile: a predictor of clinical outcome in dengue infection. Trop. Med. Int. Health TM IH 14, 576–585. https://doi.org/10.1111/j.1365-3156.2009.02261.x

van Gorp, E.C.M., Suharti, C., Mairuhu, A.T.A., Dolmans, W.M.V., van Der Ven, J., Demacker, P.N.M., van Der Meer, J.W.M., 2002. Changes in the plasma lipid profile as a potential predictor of clinical outcome in dengue hemorrhagic fever. Clin. Infect. Dis. Off. Publ. Infect. Dis. Soc. Am. 34, 1150–1153. https://doi.org/10.1086/339539

Villar, L., Dayan, G.H., Arredondo-García, J.L., Rivera, D.M., Cunha, R., Deseda, C., Reynales, H., Costa, M.S., Morales-Ramírez, J.O., Carrasquilla, G., Rey, L.C., Dietze, R., Luz, K., Rivas, E., Miranda Montoya, M.C., Cortés Supelano, M., Zambrano, B., Langevin, E., Boaz, M., Tornieporth, N., Saville, M., Noriega, F., CYD15 Study Group, 2015. Efficacy of a tetravalent dengue vaccine in children in Latin America. N. Engl. J. Med. 372, 113–123. https://doi.org/10.1056/NEJMoa1411037

Villareal, V.A., Rodgers, M.A., Costello, D.A., Yang, P.L., 2015. Targeting host lipid synthesis and metabolism to inhibit dengue and hepatitis C viruses. Antiviral Res. 124, 110–121. https://doi.org/10.1016/j.antiviral.2015.10.013

Weinglass, A.B., Kohler, M., Schulte, U., Liu, J., Nketiah, E.O., Thomas, A., Schmalhofer, W., Williams, B., Bildl, W., McMasters, D.R., Dai, K., Beers, L., McCann, M.E., Kaczorowski, G.J., Garcia, M.L., 2008. Extracellular loop C of NPC1L1 is important for binding to ezetimibe. Proc. Natl. Acad. Sci. U. S. A. 105, 11140–11145. https://doi.org/10.1073/pnas.0800936105

Welsch, S., Miller, S., Romero-Brey, I., Merz, A., Bleck, C.K.E., Walther, P., Fuller, S.D., Antony, C., Krijnse-Locker, J., Bartenschlager, R., 2009. Composition and three-dimensional architecture of the dengue virus replication and assembly sites. Cell Host Microbe 5, 365–375. https://doi.org/10.1016/j.chom.2009.03.007

Whitehorn, J., Nguyen, C.V.V., Khanh, L.P., Kien, D.T.H., Quyen, N.T.H., Tran, N.T.T., Hang, N.T., Truong, N.T., Hue Tai, L.T., Cam Huong, N.T., Nhon, V.T., Van Tram, T., Farrar, J., Wolbers, M., Simmons, C.P., Wills, B., 2016. Lovastatin for the Treatment of Adult Patients With Dengue: A Randomized, Double-Blind, Placebo-Controlled Trial. Clin. Infect. Dis. Off. Publ. Infect. Dis. Soc. Am. 62, 468–476. https://doi.org/10.1093/cid/civ949

